# Repositioning septins within the core particle

**DOI:** 10.1101/569251

**Authors:** Deborah C. Mendonça, Joci N. Macedo, Rosangela Itri, Samuel L. Guimaraes, Fernando L. Barroso da Silva, Alexandre Cassago, Richard C. Garratt, Rodrigo Portugal, Ana P. U. Araujo

## Abstract

Septins are GTP binding proteins considered to be a novel component of the cytoskeleton. They polymerize into filaments based on hetero-oligomeric core particles which, in humans, are either hexamers or octamers composed of two copies each of either three or four different septins from the 13 available. Not all combinations are possible as it is believed that these must obey substitution rules which determine that different septins must be derived from four distinct and well-established sub-groups. Here, we have purified and characterized one such combinations, SEPT5-SEPT6-SEPT7, and used TEM to derive the first structural information concerning its assembly. The full complex was purified using an affinity tag attached to only one of its components (SEPT7) and was able to bind to and perturb lipid bilayers. Although the complex assembled into elongated hexameric particles, the position of SEPT5 was incompatible with that predicted by the reported structure of SEPT2-SEPT6-SEPT7 based on the substitution rules. MBP-fusion constructs for SEPT5 and SEPT2 and immuno-staining clearly show that these septins occupy the terminal positions of the SEPT5-SEPT6-SEPT7 and SEPT2-SEPT6-SEPT7 hexamers, respectively. In so doing they expose a so-called NC interface which we show to be more susceptible to perturbation at high salt concentrations. Our results show that the true structure of the hexamer is inverted with respect to that described previously and, as such, is more compatible with that reported for yeast. Taken together, our results suggest that the mechanisms involved in spontaneous self-assembly of septin core particles and their filaments deserve further reflection.

## Introduction

Septins belong to the family of P-loop GTPases [1]. They differ from most other members in their ability to polymerize into filaments, a property which is the result of a characteristic sequence known as the Septin Unique Element [2,3]. Septins have been classically described to participate in cytokinesis, but also play roles in other important intra-cellular processes. These include acting as scaffolds in the recruitment of binding partners; as diffusion barriers in the compartmentalization of membrane proteins; in host-microorganism interaction and even in mechanotransduction [4–7].

Structurally, septins are divided into three principal domains: the N-terminal (N), GTP-binding (G) and C-terminal (C) domains. The central, G domain, is the most conserved and its ability to bind GTP is essential for the interaction between septins within filaments and is fundamental to ensure structural integrity [3,8,9]. Based on sequence similarity, the 13 mammalian septins have been subdivided into four distinct groups [10]. Representatives of either three or all four of the groups combine to form linear oligomeric core complexes, which assemble end-to-end into filaments and thence into higher-order structures [11–13].

It is well established that septin filaments can be broken down into their corresponding core complexes under conditions of high ionic strength [10,11,14]. To date, the best description of such a complex is that of the linear hexameric particle formed of two copies each of human septins 2, 6 and 7 [13]. Each septin monomer participates in two interfaces which alternate along the filament: the NC interface, involving the N and C terminal helices of the G domains; and the G interface, including the region directly involved in guanine-nucleotide-binding [3,13]. The remaining domains (particularly the coiled coil region of the C-domain) also contribute to the affinity and specificity of the NC interface [15,16].

Sirajuddin et al., 2007, using negative-stain electron microscopy and employing a version of SEPT2 fused to MBP, proposed that SEPT2 occupies the central position of the core complex and SEPT7 lies at its extremities, leading to the following arrangement for the hexamer: SEPT7-SEPT6-SEPT2-SEPT2-SEPT6-SEPT7 (which, for convenience, we will abbreviate to 7-6-2-2-6-7). This arrangement leaves a G-interface free at each end of the particle and has widely been considered to be a true description of the hexamer since the publication of the original crystal structure [13]. Subsequent studies have shown the existence of octameric core particles which incorporate SEPT9 [17–19]. In this case it has been assumed that SEPT9 occupies the terminal position by forming a G interface with SEPT7. As a consequence the octameric core particle exposes NC interfaces at its termini as is the case for yeast [20].

Kinoshita (2003) proposed that any given septin could be substituted by another from the same group, generating a physiologically viable combination. This hypothesis allows for the prediction of the position of any particular septin within the core complex and limits the number of possible combinations to 20 for hexamers and 60 for octamers. However, the generality of Kinoshita’s hypothesis is not yet fully established. One the one hand, most of the theoretically possible combinations have yet to be described *in vivo* and on the other, there have been reports of complexes which are incompatible with its basic premise [21,22]. In summary the rules governing core complex and filament assembly are far from fully established.

In order to enhance our current knowledge concerning septin heterocomplexes, we have selected a combination composed of SEPT5, SEPT6 and SEPT7 for more detailed molecular characterization. This choice was made on the basis of two criteria. Firstly, previous experiments have shown the existence of such a complex in platelets where it associates with microtubules and plays an important role in granule trafficking [23,24]. Secondly it is a conservative choice as it represents a minimal alteration when compared to the canonical SEPT2-SEPT6-SEPT7 complex. The substitution of SEPT2 by SEPT5, which belong to the same subgroup, is in accordance with Kinoshita’s predictions. Unexpectedly, we show that both of the hexameric core complexes are inverted with respect to that reported previously suggesting the need for a critical revision of the septin literature.

## Material and Methods

### Plasmid construction

cDNAs encoding full-length human SEPT2 (GenBank NM_004404), SEPT5 (GenBank NM_002688), SEPT6 (GenBank NM_145799) and SEPT7 (GenBank NM_001788), were amplified from a fetal human brain cDNA library (Clontech) by PCR and checked by automated sequencing.

Complexes (5-6-7 or 2-6-7) were co-expressed using two bi-cistronic vectors. For the first expression vector, the cDNA encoding SEPT7 (residues 29-437) was subcloned into pETDuet™-1 (NOVAGEN), using *BamH*I and *Pst*I restriction sites. SEPT7 was therefore produced in fusion with an N-terminal His-tag allowing for its purification by metal affinity chromatography, together with its interaction partners. For the second expression vector construct, the coding sequence for SEPT5 (or SEPT2) was inserted into either pRSFDuet™-1 (NOVAGEN) or pRSFMBP (details below), using *EcoR*I e *Sal*I restriction sites in both cases. Subsequently, the CDS for SEPT6 was subcloned (using *EcoR*V and *Xho*I restriction sites) into the same plasmids already containing SEPT5 (or SEPT2). The final constructs permitted the co-expression of SEPT6 and SEPT5 (or SEPT2) with or without the maltose binding protein (MBP) fused to the latter. Briefly, the pRSFMBP vector was constructed based on the plasmid pRSFDuetTM-1 (NOVAGEN) with the replacement of the coding sequence for the His-tag by that of MBP (Maltose Binding Protein) and a thrombin protease cleavage site.

### Protein expression and purification

Expression of the protein complexes was performed in *E. coli* Rosetta (DE3) as host cells. Since the protocols developed for the expression and purification of both complexes were distinct in some details, a protocol for the SEPT5-SEPT6-SEPT7 complex will be presented first, followed by the changes made in the case of the SEPT2-SEPT6-SEPT7 complex.

### SEPT5-SEPT6-SEPT7 Complex

50 mL from an overnight culture harboring both expression plasmids were inoculated into 2 L of Terrific broth medium, augmented with ampicillin (50 μg/ml), kanamycin (30 μg/ml) and chloramphenicol (34 μg/ml). Cells were grown at 37°C whilst shaking at 250 rpm. When the OD_600nm_ reached 1 to 1.2, the temperature was decreased to 20°C for 1 hour. Expression was induced with 0.25 mM IPTG (isopropyl-β-D-tiogalactopiranoside) at 20°C, 200 rpm for 16 hours. Cells were harvested by centrifugation at 6,000 × g for 40 minutes at 4°C, re-suspended in 100 ml buffer A (25 mM Tris-HCl, pH 7.8, containing 500 mM NaCl, 5 mM MgCl_2_, 5 mM β-mercaptoethanol, 5% (v/v) glycerol, with fresh addition of protease inhibitor cocktail tablets (Roche), in the presence of 0.5 mg/mL lysozyme. The suspension was incubated for 30 minutes at 4 °C. Cells were disrupted by sonication and centrifuged at 18,000 × g for 30 minutes at 4 °C. The soluble fraction was loaded onto a column containing 5 ml Ni-NTA superflow resin (Qiagen), pre-equilibrated with buffer A. The column was washed with 7 volumes of buffer A supplemented with 0.1% Triton X-100, followed by 7 volumes of buffer A. The SEPT5-SEPT6-SEPT7 complex was eluted in 50 ml of buffer A containing 400 mM imidazole.

### SEPT2-SEPT6-SEPT7 Complex

To maximize yields, SEPT2 and SEPT6 were co-expressed separately from SEPT7. Subsequently, cells were harvested by centrifugation at 5,000 × g for 10 minutes at 4°C, re-suspended and combined in 60 ml buffer B (25 mM HEPES, pH 7.8, containing 800 mM NaCl, 5 mM MgCl_2_, 1 mM β-mercaptoethanol, 5% (v/v) glycerol), in the presence of 0.1 mg/mL lysozyme. The suspension was incubated for 30 minutes at 4 °C. The cells were disrupted by sonication and centrifuged at 10,000 × g for 60 minutes at 4 °C. The soluble fraction was loaded onto a HisTrap HP 5ml column (GE Healthcare), pre-equilibrated with buffer B and then washed with 6 volumes of buffer B supplemented with 30 mM imidazole. The SEPT2-SEPT6-SEPT7 complex was eluted during a linear gradient of 25 ml of buffer B ramping from 30 to 500 mM imidazole.

After the affinity chromatography, the purified complexes were concentrated and loaded onto a Superdex 200 10/300 GL column (GE Healthcare) pre-equilibrated with buffer B. The purity and integrity of the septin complexes were analyzed on 12% SDS-PAGE. Fractions containing the complex were concentrated to 1mg / mL and aliquots of 20μL were frozen in liquid N_2_ and stored at −80 °C.

### Mass spectrometry

The individual components of the complex were identified by LC-MS/MS analysis on an ESI-micrOTOF-Q II 10234 mass spectrometer (Bruker Daltonics) and the data analyzed with MASCOT, available from http://www.matrixscience.com/search_form_select.html.

### Analysis of nucleotide content

Nucleotides were extracted from the purified protein complex samples according to the method described by Seckler et al, with some modifications [25]. Ice-cold HClO_4_ (final concentration 0.5 M) was added to the purified SEPT5-SEPT6-SEPT7 complex in buffer containing 300 mM NaCl, followed by incubation on ice for 10 min. After centrifugation at 16,000 g at 4 °C for 10 min, the supernatant was buffered and neutralized with 100 μL of KOH 3 M, 100 μL K_2_HPO_4_ 1 M, and 20 μL acetic acid to a final concentration of 0.5 M. After a centrifugation step at 16,000 g at 4 °C for 10 min, the nucleotides were analyzed using anion exchange chromatography on a Protein Pack DEAE 5 PW 7.5 mm × 7.5 cm column (Waters) driven by a Waters 2695 chromatography system. The column was equilibrated in 50mM Tris at pH 8.0 and 150 μL of each sample were loaded into the system. Elution was performed using a linear NaCl gradient (0.1–0.45 M in 10 min) at a flow rate of 1 ml/min at room temperature. The absorbance was monitored at 253 nm. As a reference, a mixture of 5 μM GDP and 5 μM GTP was used in the same buffer.

### Interaction assays with giant unilamellar vesicles (GUVs)

Phospholipids used in the GUV composition were POPC (1-hexadecanoyl-2- (9Z-octadecenoyl) -sn-glycerol-3-phosphocholine), POPS (1-palmitoyl-2-oleoyl-sn-glycerol-3-phospho-L-serine) and PIP2 (Phosphatidylinositol-4,5-bisphosphate) from Avanti Polar Lipids. The GUVs of POPC, POPC:PIP2 (97.5: 2.5 molar ratio), POPC:POPS (97.5: 2.5 molar ratio) were prepared according to the electro-formation method [26]. Interaction analyses were performed with a dilution of 100 μL of GUVs-containing sucrose buffer solution (25 mM Tris, pH 7,8, 500 mM NaCl, 5 mM MgCl_2_, 5 mM β-mercaptoetanol) (200 mMol osmolarity) freshly prepared in 600 μL of glucose-containing buffer (200 mMol osmolarity), with the subsequent addition of the SEPT5-SEPT6-SEPT7 complex to a final concentration of 26 nM. The following events were monitored by phase contrast microscopy on an Axiovert 200 (Carl Zeiss) microscope for 10 minutes and the images were recorded on a AxioCam HSm (Carl Zeiss) digital camera.

### EM and Image Processing

Purified septin complexes were diluted to 0.02 mg/ml using the final purification buffer so as to maintain the NaCl concentration between 500-800mM, adsorbed for 1 min onto glow-discharged, ultrathin carbon film supported by lacey carbon on a copper grid (Ted Pella, USA). The samples were washed in a drop of deionized water and stained in two drops of 2% uranyl acetate. Micrographs of SEPT5-SEPT6-SEPT7 complexes (with or without MBP) were recorded using an F416 camera (TVIPS, Germany) with a JEM-2100 (JEOL, Japan) microscope operated at 200 kV with a pixel size of 1.78 Å. Micrographs of MBP-SEPT2-SEPT6-SEPT7 were taken on a Talos Arctica (Thermo Fisher Scientific, USA) and were recorded using a Ceta camera with a pixel size of 2.51 Å at a defocus range of 1–3 mm underfocus. Images of high-salt (500mM) samples incubated with mouse monoclonal anti-SEPT5 were taken on a Talos F200C (Thermo Fisher Scientific, USA) microscope at 200 kV and were also recorded using a Ceta camera with a pixel size of 2.6 Å. Imagic4D software (Image Science, Germany) was used for image processing [27]. Class averages of selected particles were computed after alignment by classification and subsequent rounds of multivariate statistical analysis (MSA) using selected masks [28,29]. For better visualization of the MBP location, due to the flexible linker, one of the masks used was constructed as a rectangle on the hexameric complex, so that only information from the MBP was used for classification. For simplicity, during data processing a mask was also employed to guarantee that the MBP always appeared in the upper portion of the image. The average number of members per class was 30 particles.

### Structural Analysis

The structural characteristics of the G and NC interfaces were analyzed with the PISA server allied to visual inspection [30]. In the case of the G interface, which resides between two copies of SEPT7 within a mature filament, the recently determined crystal structure of human SEPT7 at 1.73 Å resolution (1N0B) was used. For the NC interface a chimeric model was built from two previously reported crystal structures for SEPT2 (2QNR and 2QA5). The higher resolution structure (2QNR) was used as a main model with the incorporation of the α_0_ helix (polybasic region) from 2QA5 included after structural superposition followed by structure completion and regularization with MODELLER 9.0.

Using a constant-pH Coarse Grain scheme previously devised to study protein-protein interactions, [31–33] the two interfaces were also submitted to a complexation study at pH 7 under several different salt concentrations (50, 75, 100, 150, 300, 500 and 800mM) at 298K. Titratable groups were allowed to change their protonation states according to a fast proton titration scheme shown to be accurate in reproducing experimental pKa shifts [34]. Guanosine phosphate molecules were modeled based on the guanine nucleobase titration model [35] with the inclusion of phosphates. The number of MC steps for production was at least 10^7^ after equilibration. The protein-protein radial distribution function [g(r)] was sampled with a bin size of 1 Å. From this function, the angularly averaged potential of mean force (**β**w(r)=−ln [g(r)], where **β**=1/k_B_T and k_B_ and is the Boltzmann constant) between the centers of mass of the two chains was obtained with low noise, and used to estimate the free energy of complexation (**Δ**G) under each physicochemical condition. Three independent runs were performed to assure sampling convergence.

## Results

### Characterization of the SEPT5-SEPT6-SEPT7 heterocomplex

In order to investigate the prediction of Kinoshita (2003) that SEPT5 should be able to replace SEPT2 as a component of a viable heterocomplex, we co-expressed SEPT5, SEPT6 and SEPT7 in *E. coli* Rosetta (DE3) cells. SEPT5 was produced with or without an MBP-tag and an N-terminal His-tag was added to SEPT7 to facilitate purification by metal affinity chromatography. Size exclusion chromatography proved essential for eliminating aggregates and contaminants, yielding a symmetrical peak corresponding to the stoichiometric SEPT5-SEPT6-SEPT7 complex (Fig. S1). The identity of all components was confirmed by mass spectrometry (data not shown).

### SEPT5-SEPT6-SEPT7 complex is a hexamer

Transmission electron microscopy revealed that the complex formed by SEPT5, SEPT6 and SEPT7 is a linear hexamer in which the six individual monomers can be clearly resolved (Fig 1). This result was anticipated since SEPT5 belongs to the same subgroup as SEPT2, which also forms hexamers together with SEPT6 and SEPT7. In addition, the 25 nm length of the particle agrees well with that of the SEPT2-SEPT6-SEPT7 hexamer, inferred from its crystal structure[13]. The ratio of GDP to GTP bound to this complex was 2:1 (Fig. 1B), which is consistent with that found for the 2-6-7 complex ^13^ and with the lack of catalytic activity associated with SEPT6.

**Figure 1.**
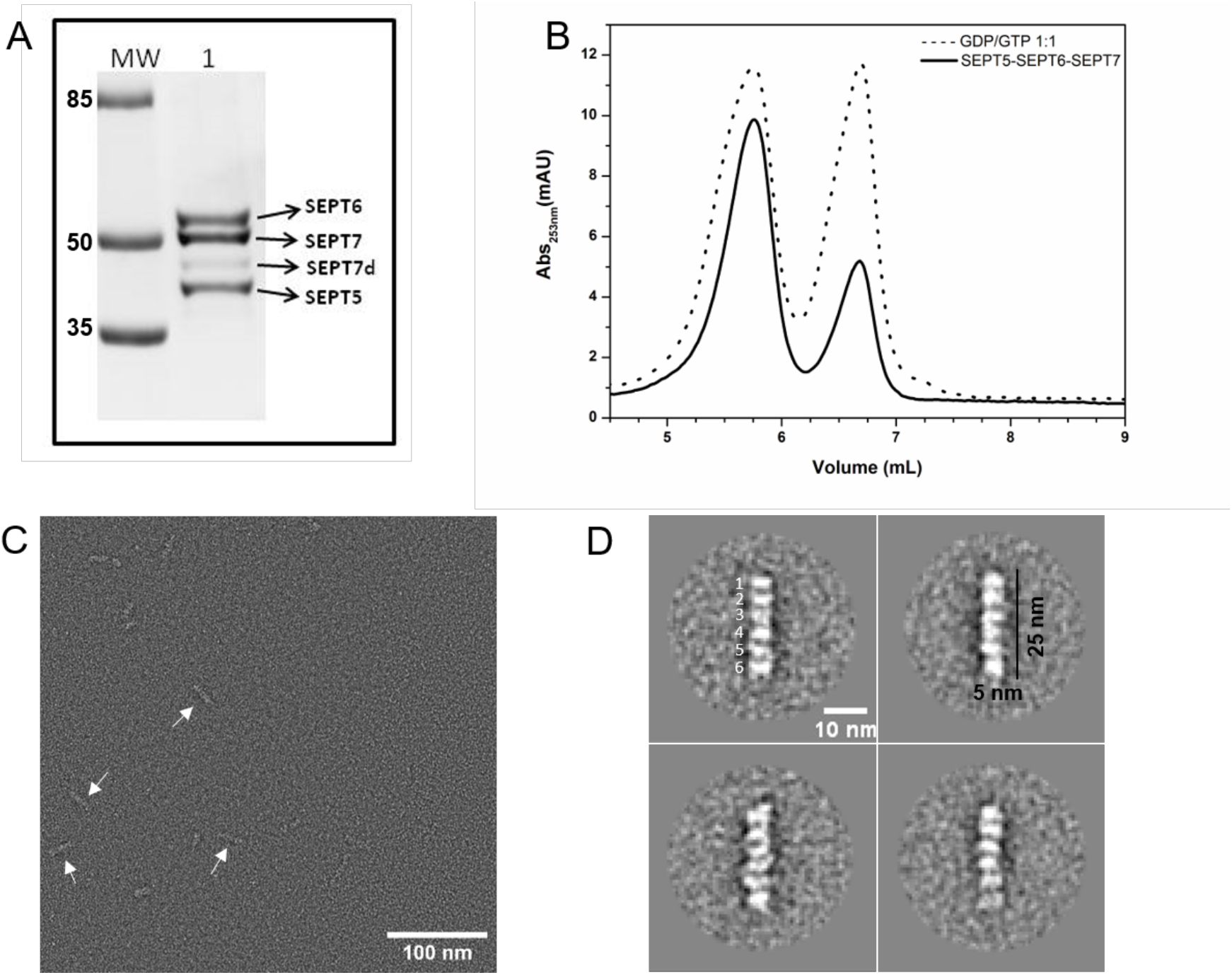
Negative-stain electron microscopy of the human SEPT5-SEPT6-SEPT7 complex. (A) 12%SDS-PAGE showing the molecular mass standard (MW in kDa) and (1) fraction corresponding to the eluted septin complex from the size exclusion chromatography. An extra, weak band (SEPT7d) corresponding to SEPT7 lacking part of the C-domain due to degradation was regularly observed during purification, similar to that described by Sirajuddin [2007] for the SEPT2-SEPT6-SEPT7 complex [36]. (B) Elution profile of guanine nucleotides from 2.4 μM of denatured purified complex. The sample was analysed by HPLC on a DEAE-5PW anion-exchange column and monitored at 253 nm. A mixture containing 5 μM of both GTP and GDP was used as a reference (dotted line); AU, absorbance units. (C) Raw electron micrograph of the purified complex at high salt concentration (800 mM). Arrows indicate hexameric particles. (D) Four representative class averages (∼30 particles each) derived from processing a total of 18,000 particles from the micrographs.

### SEPT5-SEPT6-SEPT7 complex interacts with a biomimetic membrane model

Septins typically present a polybasic region conserved in the N-domain which is responsible for lipid binding [37]. Moreover, Martínez *et al*. [2006] observed SEPT5-SEPT6-SEPT7 complexes near the plasma membrane and regulating granule fusion during platelet activation, implying direct membrane association by this complex [23].

In order to verify if the heterologous complex is functionally competent in terms of its capacity to interact with membranes, GUVs were used as biomimetic models. For all GUV compositions used, a buffer control experiment was performed to ensure that it had no effect on the vesicles. POPC was initially chosen in order to probe the interaction with the major phospholipid component in the bilayer, mimicking mammalian membranes. The addition of the SEPT5-SEPT6-SEPT7 complex to the GUVs had only a subtle effect on POPC membranes. After *circa* 5 min a limited number of small buds were observed as surface extensions to the POPC GUVs (data not shown). However, in the case of GUVs containing POPC:POPS (97.5:2.5) the complex promoted membrane shrinkage and reshaping. Figure 2 (upper panel) shows an example of such remodelling, accompanied by bud emission, over the course of about 5 minutes. A more striking result was observed with GUVs composed of POPC:PIP_2_ (97.5:2.5) where the septin complex induced dramatic modifications (Figure 2 lower panel) consistent with previous reports on the importance of phosphatidylinositol derived lipids for septin association [38,39]. A network of filamentous structures can be seen over the membrane surface on introducing the septin complex (Fig. 2 lower panel, left), followed by giant liposomal shrinkage and the rapid formation of multiple buds (Fig.2 lower panel, centre). Finally, this results in a set of smaller vesicles linked by filaments and/or small buds (lower panel, right). These observations are in accordance with Bridges *et al*. (2014) with yeast septins and confirm that the recombinant SEPT5-SEPT6-SEPT7 complex is functional in terms of membrane association and remodelling [40].

**Figure 2:**
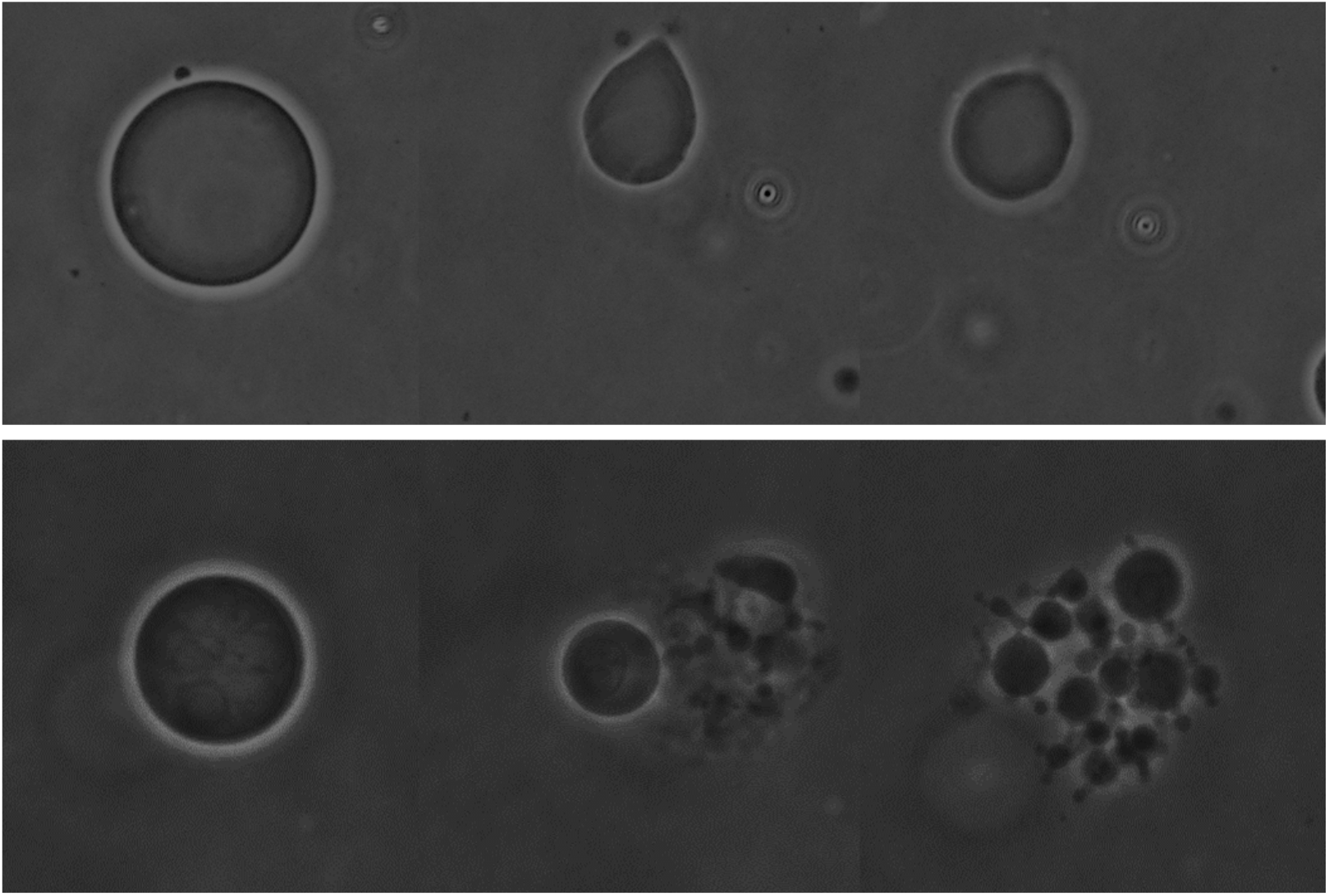
Interaction of the SEPT5-SEPT6-SEPT7 complex with GUVs. Phase contrast images of GUVs composed of POPC:POPS (upper row) and POPC:PIP_2_ (lower row) after addition of 26 nM 5-6-7 complex over the course of *circa* 5 minutes (left to right).

### SEPT5 shows an unexpected position in the hexamer

According to Kinoshita’s predictions, SEPT5 is expected to be found at the position normally occupied by SEPT2 in the canonical complex (2-6-7) since they belong to the same subgroup [10]. To test this hypothesis, TEM images of negatively stained complexes containing MBP-SEPT5 were collected and processed. Due to the flexible linker the relative position of the MBP with respect to the hexamer was expected to be variable and it was therefore essential to use a rectangular mask that would eliminate the dominant effect of the septin complex during the classification process (Fig. S2). Classes were therefore identified as a function of the position of the MBP. Different strategies were used during processing and in all cases the results showed MBP situated in variable positions centred on the end of the hexameric complex (Fig 3A). This implies that SEPT5 occupies the terminal position within the complex, different to that expected based on the currently accepted model for the canonical hexamer, 7-6-2-2-6-7 (and by implication 7-6-5-5-6-7).

**Figure 3:**
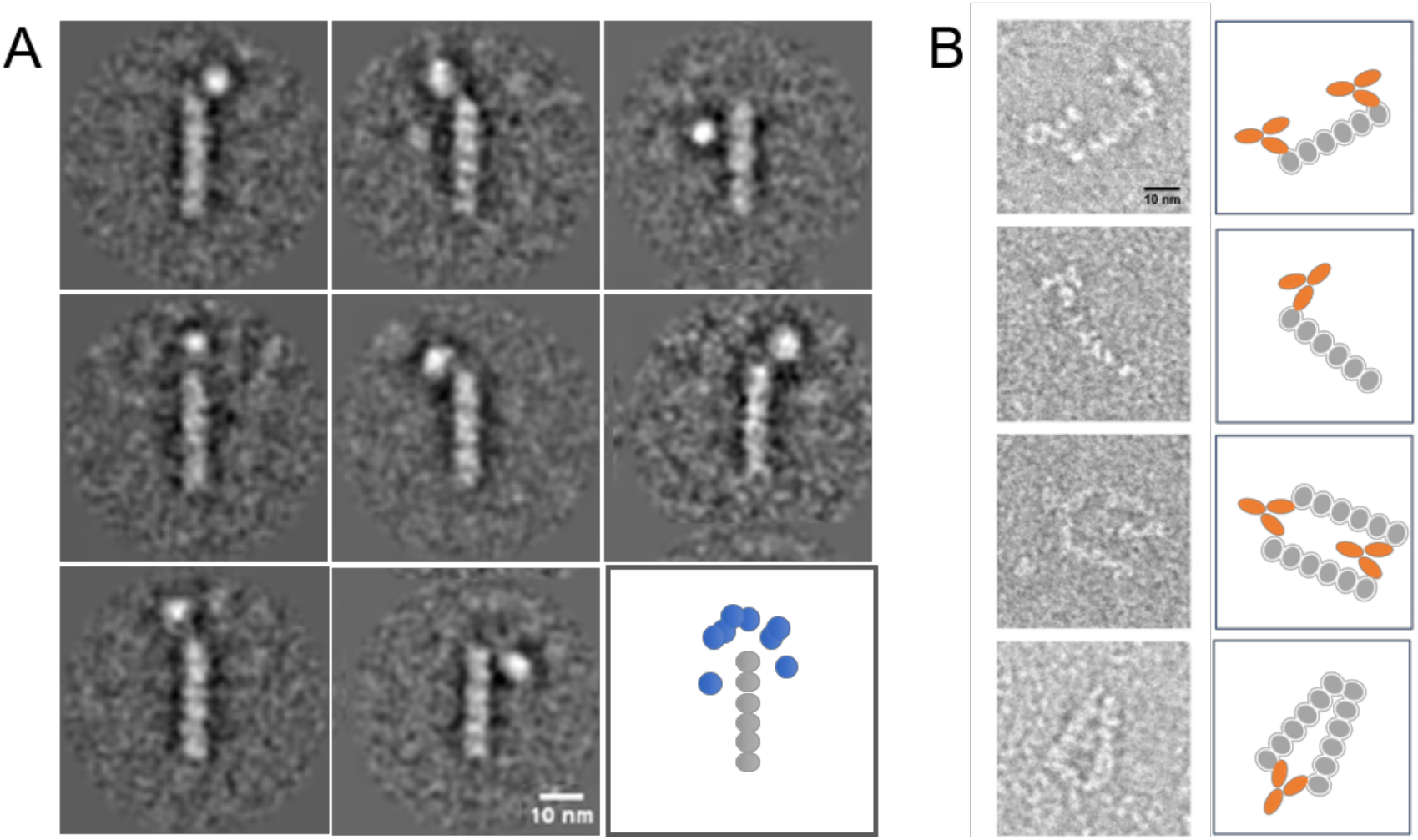
Location of MBP-SEPT5 and antibody decoration. (A) Representative class averages (∼30 particles each) of purified MBP-SEPT5-SEPT6-SEPT7 at high salt concentration (800mM). The extra density is most often positioned close to the end of the rods. On the bottom right is shown a schematic of the superimposed class averages showing the terminal monomer of the hexamer to lie at the centre of the arc defined by the different MBP positions. (B) Left, four representative raw (unaveraged) images of the sample prepared with purified human SEPT5-SEPT6-SEPT7 complex incubated with mouse anti-SEPT5 antibody at 500 mM NaCl. Right, schematic diagram of the complexes, where grey represents septin subunits and orange the antibody.

Although convincing, the unexpected nature of this result together with concerns about the flexibility of the linker between the MBP and SEPT5 led us to propose a new experiment to confirm the position of SEPT5. For this, a specific antibody was used to label SEPT5 (lacking MBP) within the complex. Examination of the raw images of particles from these micrographs confirmed the location of SEPT5 at the end of the hexamer, as shown in Figure 3B. In some cases, a single antibody is observed interacting with one end of the hexamer and in others an antibody cross-links two hexamers.

### Reviewing the position of SEPT2 in the core complex

The unexpected arrangement of the 5-6-7 hexamer suggested two possibilities; either the substitution of SEPT2 by SEPT5 leads to a genuinely different assembly or the currently accepted model is wrong. To distinguish between these possibilities, analogous TEM studies were undertaken using the canonical 2-6-7 complex, once again employing a construct in which MBP was fused to SEPT2, as per Sirajuddin (2007). Fig 4 shows eight representative class averages in which the MBP is clearly visible close to the extremity of the core complex implying that SEPT2 occupies the terminal position. The results from both complexes are therefore coherent with Kinoshita’s proposition (namely that SEPT2 and SEPT5 occupy equivalent positions) and clearly indicate that the model originally proposed is incorrect.

**Figure 4:**
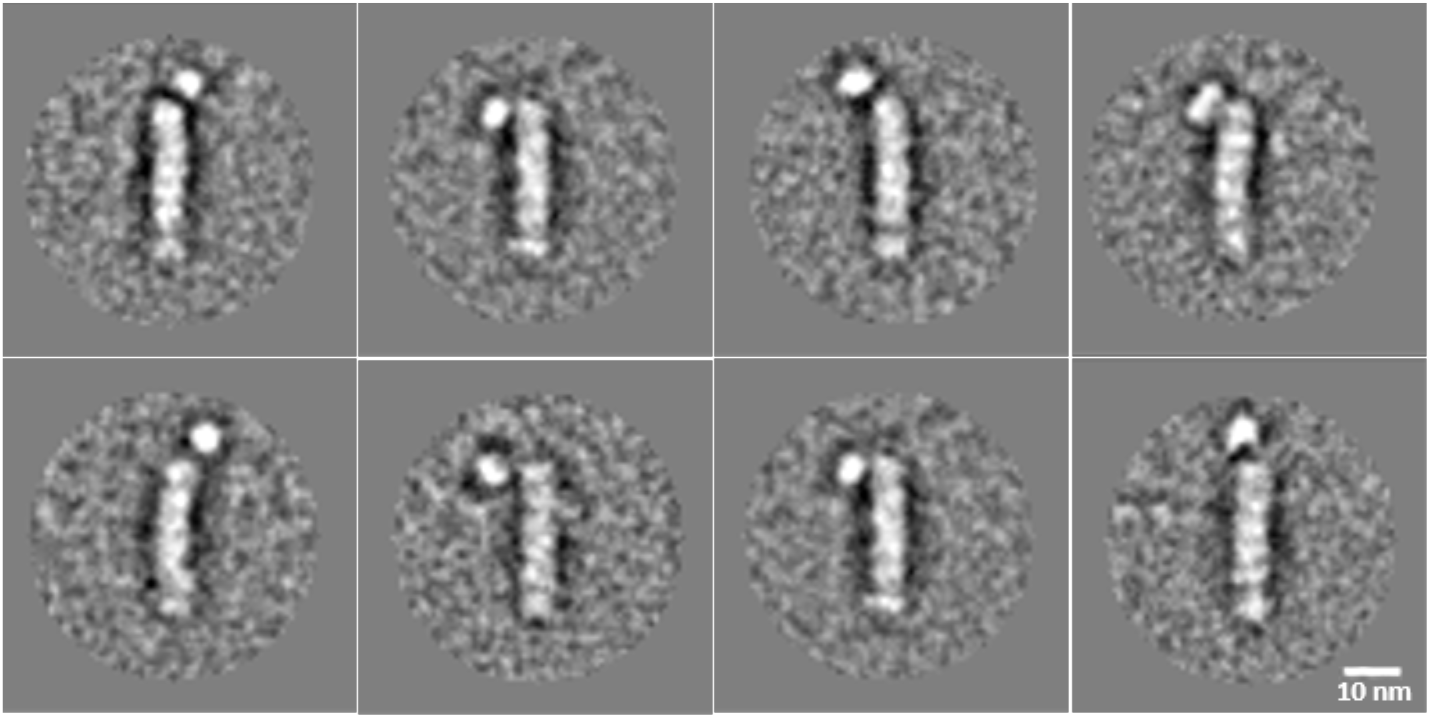
Location of SEPT2 determined by negative stain TEM using MBP-SEPT2. Representative class averages (∼30 particles each) of purified MBP2-6-7 human septin complex at high salt concentration (800mM). The extra density (MBP) is positioned close to the end of the rods, as observed for MBP5-6-7.

## Discussion

Despite recent advances, particularly at the structural level, the mechanisms underlying septin polymerization are still incompletely understood. Here we compare the organization of the individual septin components in the canonical 2-6-7 core complex with our results on 5-6-7, searching for patterns in the mode of assembly.

In the crystal structure of the human 2-6-7 complex the asymmetric unit consists of a trimer organized in the following manner; SEPT2-SEPT6-SEPT7, in which a G interface is formed between SEPT2 and SEPT6 and an NC interface between SEPT6 and SEPT7 [13]. From this trimer, the hexameric core complex (as observed with TEM under high salt conditions) could be generated by either of two crystallographic twofold axes leading to the following alternative arrangements, 7-6-2-2-6-7 or 2-6-7-7-6-2. An MBP-SEPT2 construct was used together with single particle analysis in order to define the boundaries of the particle, leading to the conclusion that SEPT2 was located at the centre of the hexamer leaving SEPT7 exposed at its extremities. Technological advances over the last decade have greatly enhanced the ease with which single particle images can be interpreted and on repeating this experiment we observe that, in fact, the hexamer is inside out with respect to that reported previously and the correct order of the subunits is therefore 2-6-7-7-6-2 (Fig. 4). Furthermore, we observe using two different approaches that this is also the case for the core particle 5-6-7-7-6-5 in accordance with Kinoshita’s proposal that septins from the same subgroup should be interchangeable [10].

One important consequence of our observation is that the interface which now lies exposed at the extremity of the core complex is an NC interface rather than G. This is the interface which is susceptible to high salt concentrations leading to depolymerization under these conditions. This interface is therefore fundamental to the polymerization process and it has been shown in yeast that it is the formation of end-to-end contacts between core complexes via lateral diffusion within the membrane which leads to polymerization [40].

The question then arises as to which of the two interfaces would be expected to more labile - the SEPT7-SEPT7 G interface or the SEPT2-SEPT2 NC interface? PISA was used to analyse both. The G interface was taken directly from the high-resolution crystal structure of the SEPT7 G-domain (pdb code 6N0B) and a model for the NC interface was generated from two incomplete crystal structures (pdb codes 2QNR and 2QA5). The two interfaces have approximately equal buried surface areas (1677 and 1651 Å^2^ per subunit respectively for NC and G) but the predicted binding energy of interaction is slightly more favourable for the G interface (−21.8 compared with −16.8 kcal/mol). The P-value, an indicator of the likelihood of a physiologically relevant interaction, is also more favourable for G than NC (0.29 compared with 0.55). Most notable, however, is that the number of charged residues involved in salt bridges at the interface is 9 at the NC interface but only 4 at the G interface. Many such interactions involve the polybasic region which is tucked into the NC interface and stabilized by compensating charges from the neighbouring subunit. At high ionic strength, with an increased dielectric constant, this interface would be expected to be destabilized exposing the SEPT2 NC surface, consistent with the new model for the hexamer.

To test this further, Monte Carlo simulations were performed and used to determine the interaction energy of the two monomers at each interface as a function of their separation. Fig. 5 shows how -βΔG (a measure of interface stability) at the optimal separation decreases as a function of the NaCl concentration. The SEPT2-SEPT2 NC interface is markedly more sensitive to the salt concentration and is clearly less stable above about 100mM. Once again this indicates that de-polymerization would be expected to occur preferentially by disruption of the SEPT2-SEPT2 NC interface rather than the SEPT7-SEPT7 G interface, consistent with exposing a SEPT2 NC interface at each end of the hexamer as we observe experimentally.

**Figure 5.**
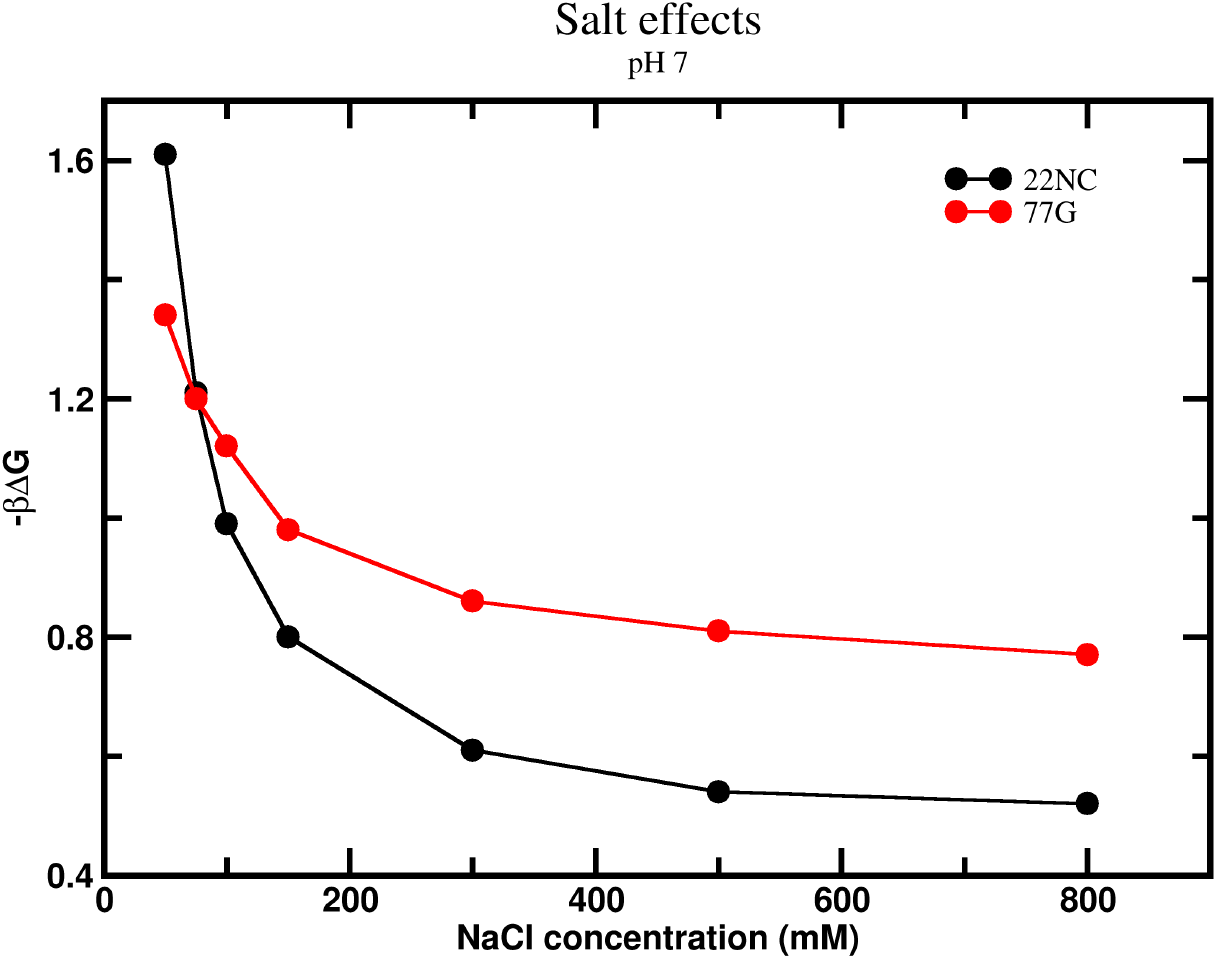
Theoretical calculations of the stability of the SEPT2-SEPT2 NC interface (black curve) and the SEPT7-SEPT7 G interface (red) as a function of NaCl concentration.

Furthermore, this proposal finds experimental support in the literature. For example, through a series of mutations, Zent *et al*. found that the G interface of SEPT7 is more stable at high salt concentrations than SEPT2. Perhaps more surprising is the fact that SEPT2 G-domains purify as homodimers which are stabilized by the G-interface rather than the physiological NC interface [13]. Once again this appears to highlight a preference for the former and the fragility of the latter.

It is pertinent to note that the octameric core complex from yeast also presents an NC interface exposed at its termini and this is similarly salt sensitive (Fig 6) [20]. In this octameric particle there is an additional septin, Cdc10. This lacks the C-terminal coiled-coil domain and is located at the center of the complex. Therefore, in both the mammalian hexamer and the yeast octamer, rupture occurs at analogous positions, namely at the NC interface which forms a homodimeric coiled coil.

**Figure 6:**
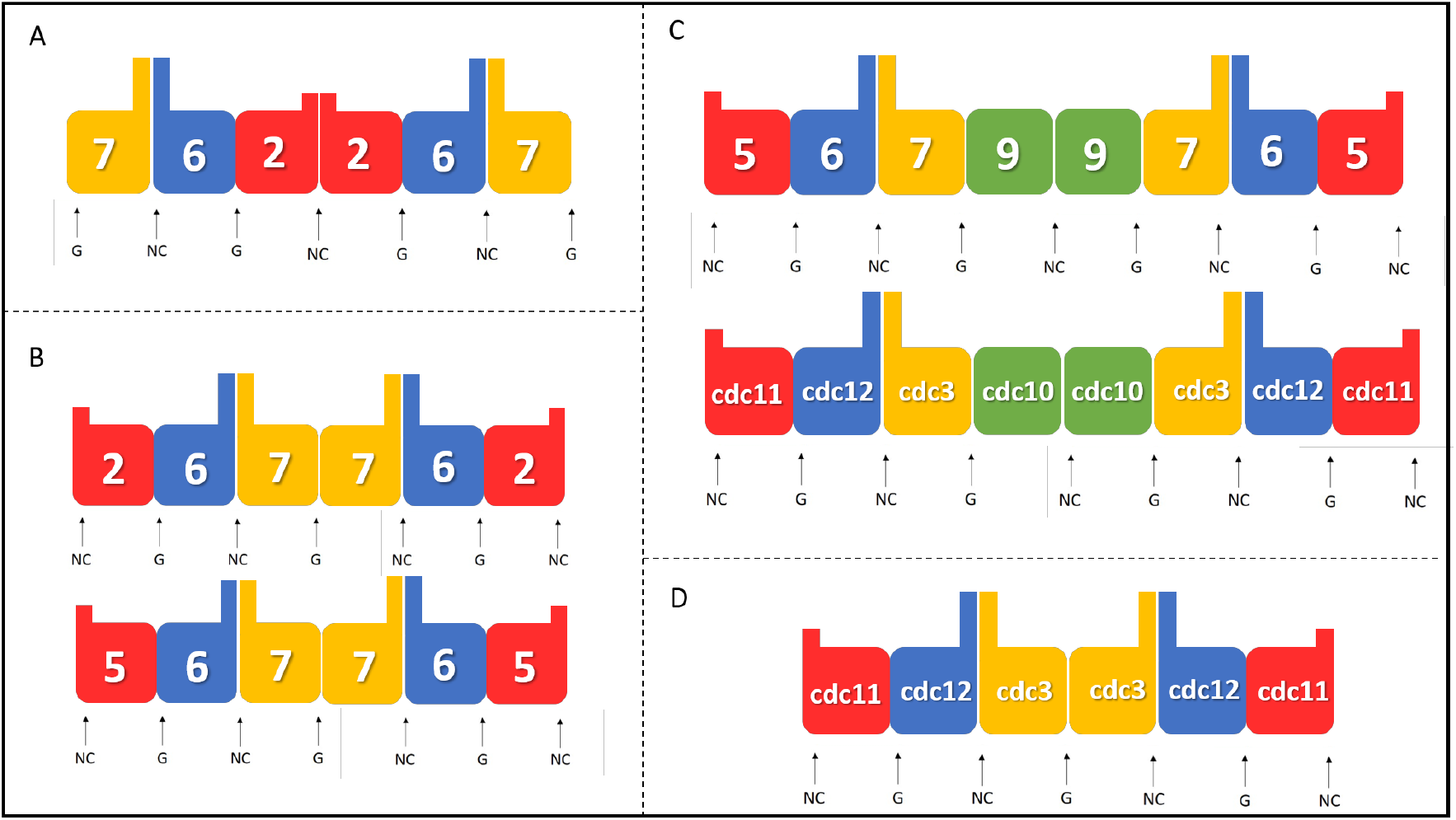
Proposition for human septin positions in the heterocomplexes. A) The canonical arrangement for the SEPT2-SEPT6-SEPT7 complex as described in the literature; B) Proposed arrangement for the 2-6-7 and 5-6-7 complexes based on the results described here; C) Proposal for the arrangement of the octameric core particle (including the SEPT9 group) and its comparison with the *S. cerevisiae* septin complex [20]; D) The hexameric particle from *S. cerevisiae* lacking cdc10 [41]. Arrows indicate NC and G interfaces.

Octameric complexes of human septins have also been reported. These include SEPT9, a member of the remaining subgroup which does not participate in the hexamer [17,19]. Cdc10 is evolutionarily closer to SEPT9 than to other mammalian septins and both lack the C-terminal coiled coil [42]. However, it has been implied that SEPT9, which interacts strongly with SEPT7 would be located at the end of the core complex [43]. This would be compatible with the standard model but not with that presented here and it is tempting to speculate that SEPT9, when present, would occupy the central position. If this were the case then the yeast and human octamers would display analogous architectures (Fig. 6C). *In vivo* studies in yeast have shown that in the absence of Cdc10, a hexamer is rescued, which has Cdc11 (analogous to the SEPT2 subgroup) at the end [41]. In this case, the central septin, Cdc3, forms a homotypic interaction via a G interface, forming a hexameric arrangement similar to the human complex proposed here (Fig 6D).

Our results change the current ideas concerning the assembly of septin filaments from their monomeric components and core complexes by placing SEPT7 at the center of the hexamer in both of the complexes studied (Fig. 6B). These appear to assemble in accordance with Kinoshita’s proposal and it is therefore reasonable to assume that our results are generic and will be applicable to other combinations of septins which have yet to be characterized experimentally [10]. More than this, our data rationalize an anomaly which was previously difficult to explain; namely that of why human septin core complexes apparently left exposed the very stable G interface whilst yeast core complexes do not. We have shown that, in fact, at high salt concentrations, they both leave NC interfaces exposed.

It is interesting to note that septin polymerization from either of the proposed core complexes (2-6-7-7-6-2 or 7-6-2-2-6-7) leads to the same sequence of monomers along the filament. Nevertheless, the way in which filaments assemble physiologically may be critically dependent on the nature of the core complex and some aspects of septin physiology may therefore need to be reconsidered in the light of the discovery reported here.

## Author Contributions

DCM, JNM, SLG planed and performed the expression and complex purifications. DCM, AC and RP performed TEM measurements and single particle analysis. JNM and RI were responsible of GUVs experiments. FLB performed molecular dynamic simulations and analyzed the data. RCG and APUA conceived and supervised the research study and secured the funding. DCM, RCG, RP and APUA wrote the manuscript, with input from the other authors.

## Acknowledgements

We thank LNNano/CNPEM for access to the EM facility via projects TEM-16717, TEM-20359 and TEM-24386. Andressa P. A. Pinto, Isabel D. Moraes, Edson Crusca, Andreza Barbosa Gomide, Humberto M. Pereira and Diego Leonardo are thanked for excellent technical assistance and Prof. William Trimble for fruitful discussions. DCM, JNM and SLG were recipients of FAPESP scholarships and APUA, RCG and RI receive fellowships from CNPq. FAPESP is thanked for financial support via thematic project 2014/15546-1 to RCG and APUA and regular project 2015/16116-3 to FLBDS.

## References

1 Weirich CS, Erzberger JP & Barral Y (2008) The septin family of GTPases: architecture and dynamics. Nat. Rev. Mol. Cell Biol. 9, 478–89.

2 Versele M, Shulewitz MJ, Cid VJ, Bahmanyar S, Chen RE, Barth P, Alber T & Thorner J (2004) Protein – Protein Interactions Governing Septin Heteropentamer Assembly and Septin Filament Organization in Saccharomyces cerevisiae. Mol. Biol. Cel 15, 4568–4583.

3 Valadares NF, Pereira H d’ M, Araujo APU & Garratt RC (2017) Septin structure and filament assembly. Biophys Rev 9, 481–500.

4 Gladfelter AS, Pringle JR & Lew DJ (2001) The septin cortex at the yeast mother – bud neck. Curr. Opin. Microbiol. 4, 681–689.

5 Caudron F & Barral Y (2009) Septins and the Lateral Compartmentalization of Eukaryotic Membranes. Dev. Cell 16, 493–506.

6 Mostowy S & Cossart P (2012) Septins: the fourth component of the cytoskeleton. Nat. Rev. Mol. Cell Biol. 13, 183–94.

7 Lam M & Calvo F (2018) Regulation of mechanotransduction : Emerging roles for septins. Cytoskeleton, 1–8.

8 Zeraik AE, Pereira HM, Santos Y V, Brandão-neto J, Spoerner M, Santos MS, Colnago LA, Garratt RC, Araújo APU & Demarco R (2014) Crystal Structure of a Schistosoma mansoni Septin Reveals the Phenomenon of Strand Slippage in Septins Dependent on the Nature of the Bound Nucleotide *. J. Biol. Chem. 289, 7799–7811.

9 Zent E & Wittinghofer A (2014) Human septin isoforms and the GDP-GTP cycle. Biol. Chem. 395, 169–180.

10 Kinoshita M (2003) Assembly of Mammalian Septins. J. Biochem. 134, 491–496.

11 Frazier JA, Wong ML, Longtine MS, Pringle JR, Mann M, Mitchison TJ & Field C (1998) Polymerization of Purified Yeast Septins: Evidence That Organized Filament Arrays May Not Be Required for Septin Function. J. Cell Biol. 143, 737–749.

12 John CM, Hite RK, Weirich CS, Daniel J, Faty M, Schla D, Kroschewski R, Winkler FK, Walz T, Barral Y & Steinmetz MO (2007) The Caenorhabditis elegans septin complex is nonpolar. EMBO J. 26, 3296–3307.

13 Sirajuddin M, Farkasovsky M, Hauer F, Kuhlmann D, Macara IG, Weyand M, Stark H & Wittinghofer A (2007) Structural insight into filament formation by mammalian septins. Nature 449, 311–117.

14 Field CM, Al-Awar O, Rosenblatt J, Wong ML, Alberts B & Mitchison TJ (1996) A Purified Drosophila Septin Complex Forms Filaments and Exhibits GTPase Activity. J. Cell Biol. 133, 605–616.

15 Marques I de A, Valadares NF, Garcia W, Damalio JCP, Macedo JNA, de Araújo APU, Botello CA, Andreu JM & Garratt RC (2012) Septin C-Terminal Domain Interactions: Implications for Filament Stability and Assembly. Cell Biochem. Biophys. 62, 317–328.

16 Sala FA, Valadares NF, Macedo JNA, Borges JC & Garratt RC (2016) Heterotypic Coiled-Coil Formation is Essential for the Correct Assembly of the Septin Heterofilament. Biophys. J. 111, 2608–2619.

17 Sellin ME, Sandblad L, Stenmark S, Gullberg M & Kellogg DR (2011) Deciphering the rules governing assembly order of mammalian septin complexes.

18 Sandrock K, Bartsch I, Bläser S, Busse A, Busse E & Zieger B (2011) Characterization of human septin interactions. Biol. Chem. 392, 751–761.

19 Kim MS, Froese CD, Estey MP & Trimble WS (2011) SEPT9 occupies the terminal positions in septin octamers and mediates polymerization-dependent functions in abscission. J. Cell Biol. 195, 815–826.

20 Bertin A, Mcmurray MA, Grob P, Park S, Iii GG, Patanwala I, Ng H, Alber T, Thorner J & Nogales E (2008) Saccharomyces cerevisiae septins : Supramolecular organization of heterooligomers and the mechanism of filament assembly. PNAS 105.

21 Nagata K, Asano T, Nozawa Y & Inagaki M (2004) Biochemical and Cell Biological Analyses of a Mammalian Septin Complex, Sept7 / 9b / 11 *. J. Biol. Chem. 279, 55895–55904.

22 Lukoyanova N, Baldwin SA & Trinick J (2008) 3D Reconstruction of Mammalian Septin Filaments., 1–7.

23 Martinez C, Corral J, Dent JA, Sesma L, Vicente V & Ware J (2006) Platelet septin complexes form rings and associate with the microtubular network. J. Thromb. Haemost. 4, 1388–1395.

24 Neubauer K & Zieger B (2017) The Mammalian Septin Interactome. Front. Cell Dev. Biol. 5, 1–9.

25 Seckler R, Wu GM & Timasheff SN (1990) Interactions of tubulin with guanylyl- (beta-gamma-methylene)diphosphonate. formation and assembly of a stoichiometric complex. J. Biol. Chem. 265, 7655–7661.

26 Angelova MI & Dimitrov DS (1986) Liposome Electroformation. Faraday Discuss 81, 303–311.

27 Heel M van, Portugal R V., Rohou A, Linnemayr C, Bebeacua C, Schmidt R, Grant T & Schatz M (2012) Four-dimensional cryo-electron microscopy at quasi-atomic resolution: IMAGIC 4D. In International Tables for Cristallography (E. Arnold, D. M. Himmel MGR, ed), Second, pp. 624–628. John Wiley & Sons, Inc.

28 Afanasyev P, Seer-Linnemayr C, Ravelli RBG, Matadeen R, De Carlo S, Alewijnse B, Portugal R V., Pannu NS, Schatz M & van Heel M (2017) Single-particle cryo-EM using alignment by classification (ABC): the structure of *Lumbricus terrestris* haemoglobin. IUCrJ 4, 678–694.

29 Heel M Van, Portugal RV & Schatz M (2016) Multivariate Statistical Analysis of Large Datasets : Single Particle Electron. Open J. Stat. 6, 701–739.

30 Krissinel E & Henrick K (2007) Inference of Macromolecular Assemblies from Crystalline State. J. Mol. Biol. 372, 774–797.

31 Forsman J, Chatterton DEW, Åkesson T, Persson BA & Lund M (2010) Molecular evidence of stereo-specific lactoferrin dimers in solution. Biophys. Chem. 151, 187–189.

32 Delboni LA & Barroso da Silva FL (2016) On the complexation of whey proteins. Food Hydrocoll. 55, 89–99.

33 Barroso Da Silva FL, Pasquali S, Derreumaux P & Dias LG (2016) Electrostatics analysis of the mutational and pH effects of the N-terminal domain self-association of the major ampullate spidroin. Soft Matter 12, 5600–5612.

34 Barroso da Silva FL & Mackernan D (2017) Benchmarking a Fast Proton Titration Scheme in Implicit Solvent for Biomolecular Simulations. J. Chem. Theory Comput. 13, 2915–2929.

35 Barroso da Silva FL, Derreumaux P & Pasquali S (2018) Protein-RNA complexation driven by the charge regulation mechanism. Biochem. Biophys. Res. Commun. 498, 264–273.

36 Sirajuddin M (2007) Structural studies on mammalian septins – New insights into filament formation.

37 Zhang J, Kong C, Xie H, McPherson PS, Grinstein S & Trimble WS (1999) Phosphatidylinositol polyphosphate binding to the mammalian septin H5 is modulated by GTP. Curr. Biol. 9, 1458–1467.

38 Tanaka-takiguchi Y & Kinoshita M (2009) Report Septin-Mediated Uniform Bracing of Phospholipid Membranes. Curr. Biol. 19, 140–145.

39 Beber A, Alqabandi M, Prévost C, Viars F, Lévy D, Bassereau P, Bertin A & Mangenot S (2018) Septin-based readout of PI (4, 5) P2 incorporation into membranes of giant unilamellar vesicles. Cytoskeleton, 1–12.

40 Bridges AA, Zhang H, Mehta SB, Occhipinti P & Tani T (2014) Septin assemblies form by diffusion-driven annealing on membranes. PNAS 111, 2146–2151.

41 Mcmurray MA, Bertin A, Iii GG, Lam L, Nogales E & Thorner J (2012) Septin Filament Formation is Essential in Budding Yeast. Dev. Cell 20, 540–549.

42 Pan F, Malmberg RL & Momany M (2007) Analysis of septins across kingdoms reveals orthology and new motifs. BMC Evol. Biol. 7, 1–17.

43 Nakahira M, Macedo JNA, Seraphim TV, Cavalcante N, Souza TACB, Damalio JCP, Reyes LF, Assmann EM, Alborghetti MR, Garratt RC, Araujo APU, Zanchin NIT, Barbosa JARG & Kobarg J (2010) A Draft of the Human Septin Interactome. PLoS One 5.

