## Supplemental Material for "Repositioning septins within the core particle"

**Supplementary material**

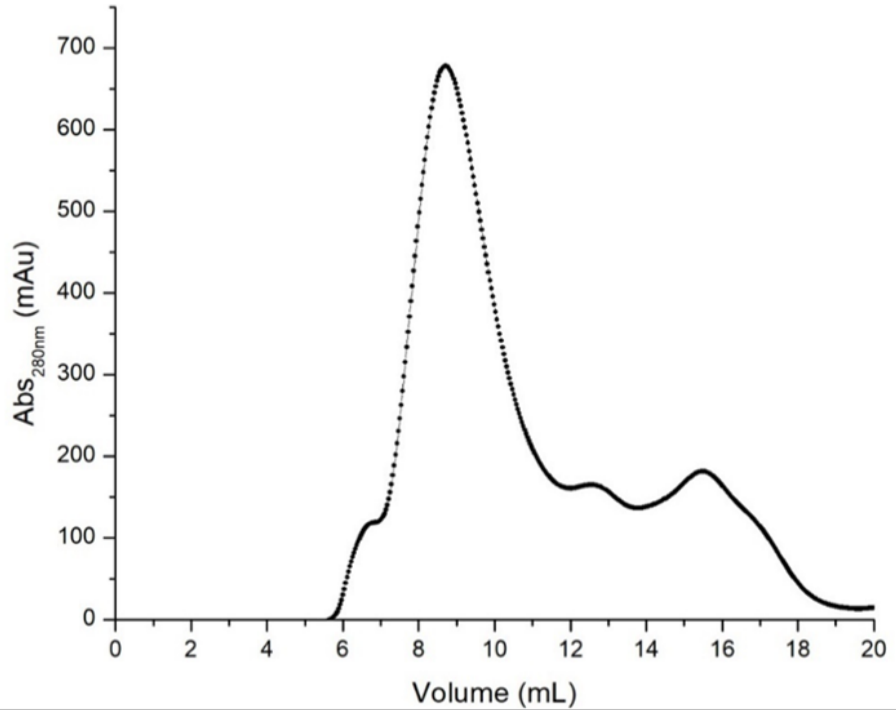


**Figure S1:** Elution profile of the SEPT5-SEPT6-SEPT7 complex from a molecular exclusion column Superdex 200. The fraction collected from 8-9 mL was used for immunolabeling and negative stain analysis.
